# Mosaicism of BRAF(V600E) in Healthy Skin Predicts Neurodegeneration in Histiocytosis

**DOI:** 10.1101/2025.11.17.688335

**Authors:** Jean-François Emile, Zofia Hélias-Rodzewicz, Matthias Papo, Irena Antonia Ungureanu, Chooyoung Baek, Félicien Triboulet, Tsz Hung Wong, Rim Ben Jannet, Nathalie Terrones, Sihame Es-Qalli, Paul-Marie Suret, Baptiste Reig, Alain Atallah, Anna Semenov, Raphael Degrave, Marianne Nguyen, Riccardo Sutera, Elina Siliogka, Pierre-Louis Cariou, Grégoire Prevost, Jeanne de la Rochefoucauld, Homa Adle-Biassette, Isabelle Plu, Alexis Mathian, Cyril Catelain, Jerome Razanamahery, Claire Ewenczyk, Sébastien Héritier, Fleur Cohen-Aubart, Pierre Hirsch, Zahir Amoura, Jean Donadieu, Ahmed Idbaih, Julien Haroche

## Abstract

Most histiocytoses harbor an oncogenic somatic alteration activating the MAP kinase pathway, the most frequent of which is *BRAF^V600E^*. Targeted therapy with BRAF or MEK inhibitors is highly effective, but neurodegeneration remains a severe complication. Recent data suggest that neurodegeneration may be secondary to mosaicism for *BRAF^V600E^* in microglia. In the prospective observational TARGET-HISTIO cohort study, we investigated extra-CNS mosaicism of *BRAF^V600E^* in a cohort of 124 adults with histiocytosis. Seeking widespread mosaicism, we analyzed healthy skin biopsies from areas without histiocytosis or melanocytic infiltration using high-sensitivity digital polymerase chain reaction. *BRAF^V600E^*was detected in healthy skin in 57/67 (85%) patients with *BRAF^V600E^*histiocytosis and was present in all second and third skin biopsies (n=6). *BRAF^V600E^*was absent from skin biopsies in all 37 patients with histiocytosis harboring another oncogenic mutation. *BRAF^V600E^* was also present in skin biopsies of four /20 patients with histiocytosis of unknown molecular status. Cells harboring *BRAF^V600E^* share characteristics with resident dermis macrophages. Only 58% of patients with *BRAF^V600E^* in skin also had *BRAF^V600E^* in blood leukocytes. Variant allele frequency in skin and blood leukocytes was 16 times lower than in histiocytosis, and frequency in skin was independent of treatment with BRAF/MEK inhibitors. None of the patients with wild type *BRAF* alleles in skin had neurodegeneration, whereas 22/61 (36%) of patients with *BRAF^V600E^* in skin had neurodegeneration. This study demonstrates that mosaicism of *BRAF^V600E^*in healthy skin is frequent in patients with histiocytosis and is strongly associated with neurodegeneration.

**Key Points:** Low allele frequency of the oncogene BRAF V600E is frequent in healthy skin of patients with histiocytosis.

Patients without BRAF V600E in skin do not develop neurodegeneration, while 36% of those with BRAF V600E mosaicism do.

## INTRODUCTION

Histiocytoses comprise a group of rare diseases characterized by tissue infiltration by cells harboring macrophage or dendritic cell markers.^1^ Histiocytoses can occur at any age and involve any tissues or organs, leading to highly heterogeneous clinical symptoms and frequently a delayed diagnosis. The most frequent subtypes of histiocytosis are Langerhans cell histiocytosis, Erdheim Chester disease, Rosai-Dorfman-Destombes disease, and xanthogranuloma, some of which can also coexist. Most histiocytoses subtypes are clonal neoplasms with oncogenic somatic alterations of genes activating the MAP kinase cell-signaling pathway.^2^ The most common mutation is *BRAF^V600E^*, present in 50–60% of patients with Langerhans cell histiocytosis and Erdheim Chester disease,^3,4^ but more rarely in other subtypes. *BRAF^V600E^*involves histiocytes but has also been detected in blood leukocytes and bone marrow progenitors.^5,6^

The discovery of *BRAF^V600E^* has led to use of BRAF inhibitors,^4,7,8^ and therapies targeting the MAP kinase pathway have rapidly emerged as the gold standard for patients with life-threatening or organ-damaging histiocytosis.^9,10^ Indeed, most patients benefit from a rapid and prolonged tumor response with BRAF, MEK, ALK, RET, and/or NTRK inhibitors.^2,4,7,8,11–13^ Histiocytoses can infiltrate the central nervous system, as well as any other tissues. Some patients with Langerhans cell histiocytosis or Erdheim Chester disease can also have symptoms unrelated to tumors but considered as neurodegeneration. The hallmark of neurodegeneration is cerebellar ataxia, which can associate with cognitive dysfunction and behavioral disturbances. On brain magnetic resonance imaging (MRI), neurodegeneration appears as non-expansile, T2, and fluid-attenuated inversion recovery (FLAIR) intense lesions in the cerebellar peduncles, medial cerebellar structures, and/or basal ganglia.^14^

The mechanism of neurodegeneration is not well understood. Interestingly, Geissmann’s team^15^ described a model of mosaic mice that exhibited targeted expression of *BRAF^V600E^* within macrophages of the brain parenchyma (microglia). These mice developed neurodegeneration that was very similar to that observed in patients with histiocytosis, and the symptoms were delayed by treatment with BRAF inhibitors. Recently, the same group reported micro-mosaicism of *BRAF^V600E^* in microglia in eight patients with Langerhans cell histiocytosis or Erdheim Chester disease, half of whom had clinical symptoms of neurodegeneration.^16^ The toxicity of microglia on neurons is further supported by cocultures of macrophages and neurons derived from *BRAF^V600E^*-induced pluripotent stem cells of patients with histiocytosis.^17^

We investigated extra-cerebral mosaicism of *BRAF^V600E^*in a large cohort of patients with histiocytosis in France. Our study focused on skin, because this tissue is easily available for histology and molecular analysis.

## METHODS

The present work is an ancillary study of the prospective observational cohort study TARGET-HISTIO (ClinicalTrials.gov number, NCT04437381). The diagnosis of histiocytosis was established according to international criteria^18^ and was confirmed by the French histiocytosis board. The study was approved by the French Ethics and Scientific Committee for Health Research, Studies and Evaluations (CESREES 2814848 bis) and was conducted in accordance with the Declaration of Helsinki. Patients provided written informed consent. Lumbar skin biopsy was considered minimally invasive according to French regulations.

Histiocytosis involvement of the central nervous system was based on clinical examination and review of the clinical records available at the time of the lumbar skin biopsy; identification was thus blinded to the mutational status of the skin. Diagnosis of neurodegeneration was based primarily on clinical and/or MRI signs of cerebellar ataxia.^14^ Head MRIs of a subgroup were reviewed blindly of the results from healthy skin.

Samples were collected by Ambroise Paré BioBank (ISO 20387). The inclusion of patients started in October 2024. Samples of healthy skin, from the margins of resected basal cell or epidermoid carcinomas, were obtained retrospectively in five patients. The other healthy skin samples were obtained prospectively: punch biopsies (three mm in diameter) were taken from clinically healthy skin in the lumbar region after the administration of local analgesia. For the first 31 patients the skin biopsies were fixed in formalin, while all the other biopsies (collected from January to May 2025) were frozen.

All skin biopsies underwent histology control by a pathologist (J.F.E.) before DNA extraction. Fifteen fixed skin biopsies were also stained with CD1a (clone O10) and SOX10 (clone EP268). DNA extraction was performed using QIAamp DNA mini (Qiagen) for frozen skin, RSC DNA FFPE (Maxwell) for fixed skin, and RSC whole blood DNA kit (Maxwell) for blood leukocytes, according to manufacturer’s instructions. Digital polymerase chain reaction of *BRAF^V600E^* and control wild type *BRAF* simultaneous quantifications (ISO 15189) was performed as described previously.^19^ Negative samples with less than 10,000 wild type *BRAF* copies were excluded.

The location of mutant cells was investigated by *BRAF^V600E^*-specific in-situ hybridization on frozen healthy skin biopsies. This procedure was conducted with a single-plex BaseScope Probe (Bio-Techne) specific to *BRAF^V600E^* and *BRAF* wild type alleles and was performed on frozen tissue samples according to manufacturer’s instructions. Immunohistochemistry was performed using CD163 (clone 10D6) and CD4 (clone 4B12).

## RESULTS

### STUDY POPULATION

Median age of the 124 patients was 61 years (range 19.1 to 88.8 years), with 49 females and 75 males (Table S1 in the Supplementary Appendix). Most patients had Erdheim-Chester disease (n=68), Langerhans cell histiocytosis (n=21) or both of these diseases (n=11) (Fig. 1). An oncogenic mutation activating the MAP kinase cell-signaling pathway was identified in histiocytosis-infiltrated biopsies from 104 (84%) patients, and corresponded to *BRAF^V600E^* in 67 (64%) and *MAP2K1* in 27 (26%). Neurodegeneration was diagnosed in 22 (18%) patients.

**Figure 1.**
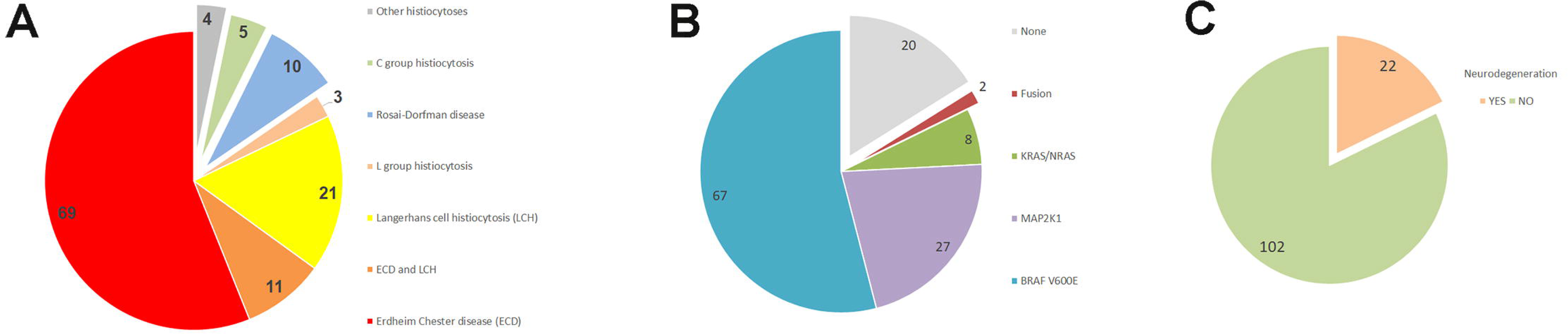
Study population. Type of histiocytosis (A) and molecular status (B) of the 124 patients. Frequency of neurodegeneration of these patients (C).

### SKIN BIOPSY RESULTS

Histological analysis of skin sections excluded the presence of histiocytosis and melanocytic tumor infiltration in all samples, and confirmed the presence in all cases of both dermis and epidermis within the sections submitted for molecular analysis. Additional staining performed in 15 formalin-fixed samples confirmed the presence of healthy intraepidermal CD207-positive Langerhans cells and SOX10-positive melanocytes, and the absence of nevi.

Healthy skin samples of 61 patients contained the oncogenic variant *BRAF^V600E^*. The variant allele frequency in positive cases was low, with a median of 0.29% (range, 0.01 to 3.08%). The number of wild type *BRAF* copies in negative skin samples was high, with a median of 17,184 (range, 13,302 to 25,419).

For six patients with *BRAF^V600E^*-positive skin, two (n=5) or three (n=1) healthy skin biopsies were available for analysis. All these additional biopsies also contained *BRAF^V600E^*, suggesting that the oncogenic variant was disseminated throughout the skin body surface.

### CORRELATION OF *BRAF^V600E^* IN HEALTHY SKIN WITH HISTIOCYTOSIS CHARACTERISTICS

The mutational status of histiocytoses was previously obtained by molecular analysis of histiocytosis-infiltrated biopsies (Fig. 1). *BRAF^V600E^*was detected within healthy skin in 85.1% of patients with *BRAF^V600E^*histiocytosis. By contrast, *BRAF^V600E^* was undetectable in all healthy skin biopsies (n=37) of patients with histiocytosis harboring another oncogenic mutation (Table S2 in the Supplementary Appendix). The mutational status of histiocytosis of 20 patients was unknown; *BRAF^V600E^*was detected in the healthy skin of four of these individuals.

### *BRAF^V600E^* IN OTHER SAMPLES FROM THE PATIENTS

The variant allele frequency of *BRAF^V600E^* was significantly higher within histiocytosis-infiltrated biopsies than within healthy skin biopsies (*P*=0.000002, Student’s test). Median variant allele frequency in healthy skin was 16 times lower than in samples with histiocytosis (0.29 vs. 4.69) (Fig. 2A).

**Figure 2.**
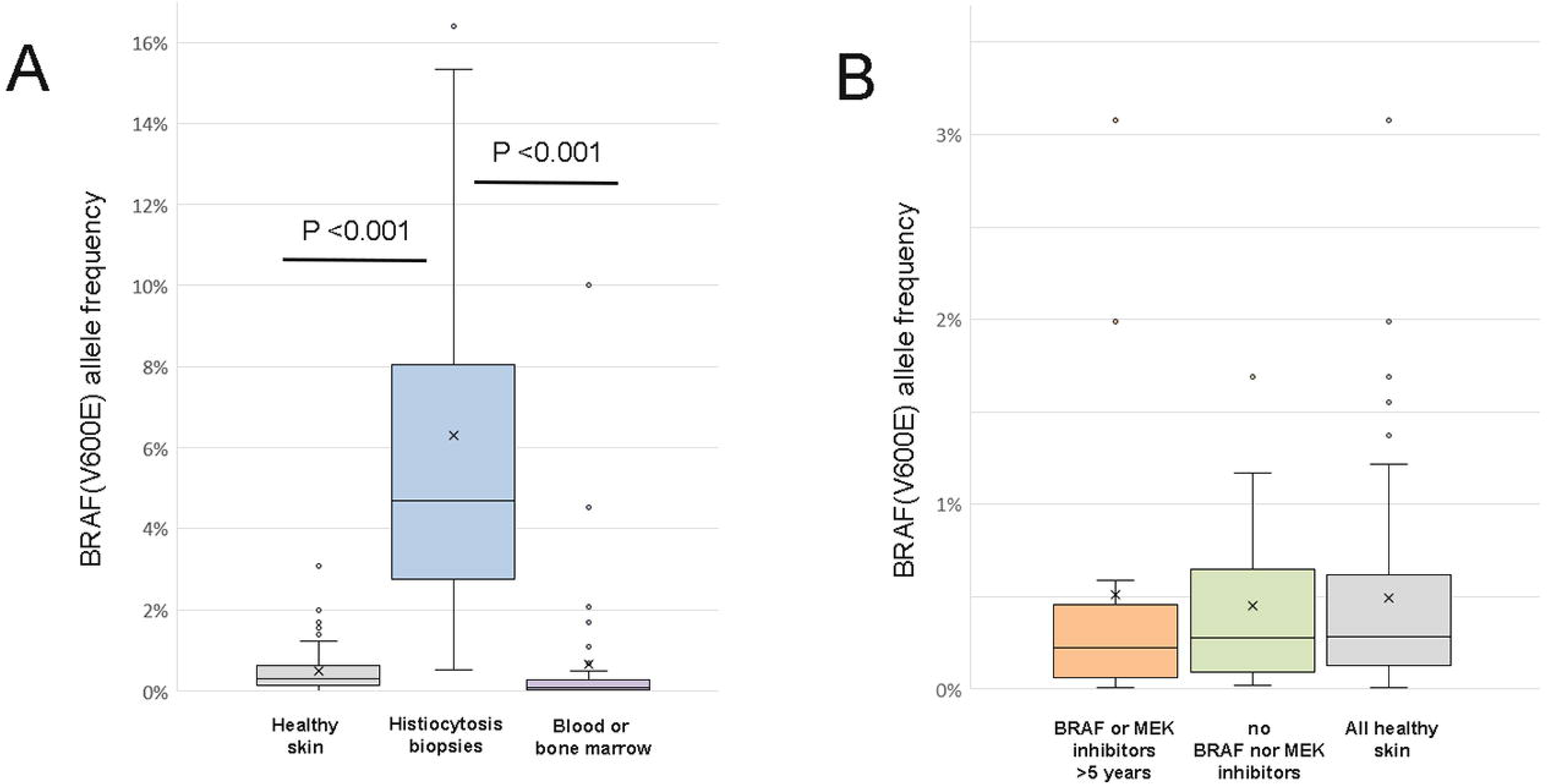
Detection of BRAFV600E Within Healthy Skin or Histiocytosis Biopsies, or Blood/Bone Marrow Cells. Variant allele frequencies were significantly lower in healthy skin and blood/bone marrow cells than in histiocytosis-infiltrated biopsies (A). Variant allele frequency of BRAFV600E in skin was not related to long-term treatment of patients with BRAF/MEK inhibitors (B).

Blood leukocytes were collected on the same day as healthy skin biopsies, whereas bone marrow DNA was obtained from samples collected up to five years earlier. There was a good correlation between the presence of *BRAF^V600E^*in blood and bone marrow cells (Table S3 in the Supplementary Appendix). As in healthy skin, median variant allele frequencies were low (0.04 in blood and 0.14 in bone marrow) (Fig. 2A), and median number of wild type *BRAF* copies of negative samples was high (20,647 in blood and 18,489 in bone marrow). Blood and/or bone marrow cells were wild type in 41.7% of patients with *BRAF^V600E^*-positive healthy skin.

### *BRAF^V600E^* IN HEALTHY SKIN ON BRAF/MEK INHIBITOR TREATMENT

Several patients (41/61, 67.2%) with *BRAF^V600E^*-positive healthy skin were treated with BRAF and/or MEK inhibitors before skin biopsy. Due to adaptation to toxicities or disease activity, and/or patient choice, almost all had had changes to the dosage, type (vemurafenib, dabrafenib, encorafenib, cobimetinib, trametinib and binimetinib), and combination (monotherapy or bitherapy) of these drugs during follow-up. However, 16 patients were on treatment with BRAF and/or MEK inhibitors for longer than 5 years at the time the healthy skin was biopsied. The variant allele frequency of *BRAF^V600E^* in these patients did not differ from that of all positive cases (Fig. 2B) or from the patients never treated with BRAF/MEK inhibitors.

### IDENTIFICATION OF SKIN CELLS HARBORING *BRAF^V600E^*

On *BRAF^V600E^*-specific in-situ hybridization, mutant cells were detected exclusively within the dermis and none in the epidermis. They were primarily found surrounding skin adnexa or blood vessels (Fig. 3). *BRAF^V600E^* cells colocalized with macrophage markers such as CD163 (serial sections) and CD4 (double staining).

**Figure 3.**
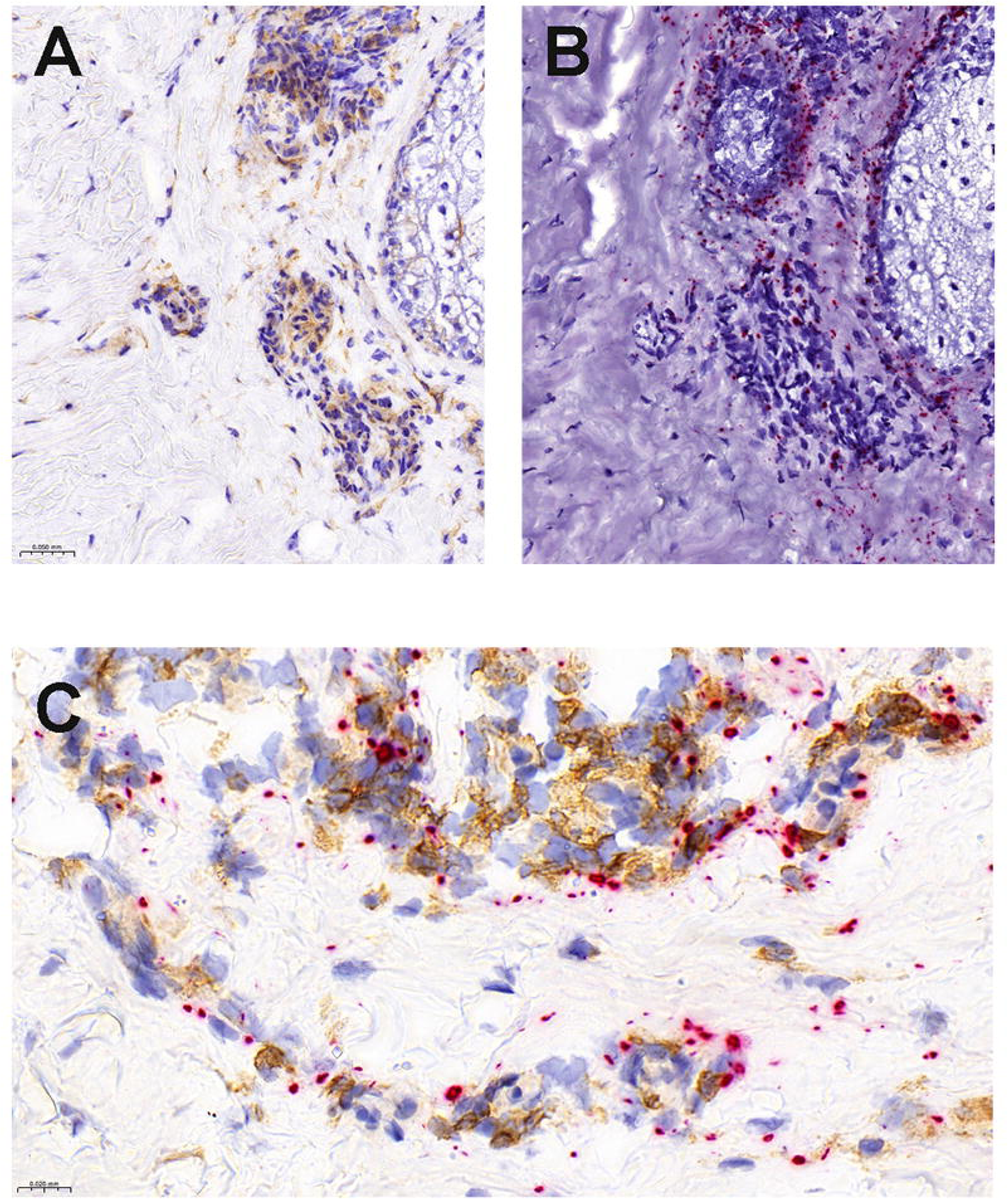
Detection of RNA Coding for BRAFV600E using Chromogenic In-Situ Hybridization. Serial sections of a healthy skin biopsy, stained with CD163 (A, brown staining) and with BRAFV600E in-situ hybridization showing red dots within the dermis, around dermal adnexa and blood vessels (B) (original magnification 220); Double staining with in-situ hybridization (red dots) and CD4 (brown staining) (original magnification 600).

### PREDICTIVE VALUE OF NEURODEGENERATIVE CEREBELLAR ATAXIA

Among the 124 patients with confirmed histiocytosis, none with wild type skin (n=63) had clinical sign of cerebellar ataxia. By contrast, 22/61 (36%) of patients with *BRAF^V600E^*-positive healthy skin had cerebellar ataxia Thus, testing of healthy skin for *BRAF^V600E^* had 100% negative predictive value for cerebellar ataxia.

Review of head MRI, blinded to healthy skin status, was performed in a subset of 38 patients. Twenty-one patients were free of cerebellar involvement both clinically and on MRI. Fourteen patients had both clinical and MRI evidence of cerebellar involvement, all of whom had *BRAF^V600E^*-positive skin. The last three had cerebellar imaging suggestive of cerebellar involvement but without ataxia reported in their clinical records; two were *BRAF^V600E^*-positive, while one was wild type on skin.

In most cases, MRI lesions were suggestive of histiocytosis-related neurodegeneration (Fig. 4A), while a few patients had atypical lesions. A 33-year-old woman presented with severe clinical ataxia and major atrophy of the cerebellum on MRI (Fig. 4B). Anamnesis revealed that she was diagnosed with Langerhans cell histiocytosis at age 23, with diabetes insipidus and typical lung involvement on computed tomography scan, without any other localization. As histiocytosis was not active at this time, biopsy was not performed, and no specific treatment was proposed. She has *BRAF^V600E^*-positive skin, and high-dose BRAF inhibitor treatment has now been offered. A 29-year-old man also presented with severe clinical cerebellar ataxia, and had both tumor and neurodegenerative lesions on brain MRI (Fig. 4C). Although *BRAF^V600E^* was detected in his brain biopsy, histology failed to confirm histiocytosis infiltration. He had three skin biopsies showing *BRAF^V600E^*. The tumor board stated a diagnosis of L group histiocytosis, and he had partial clinical and MRI responses after 3 months of treatment combining dabrafenib (100 mg twice daily) and trametinib (1.5 mg once daily) (Fig. 4C).

**Figure 4.**
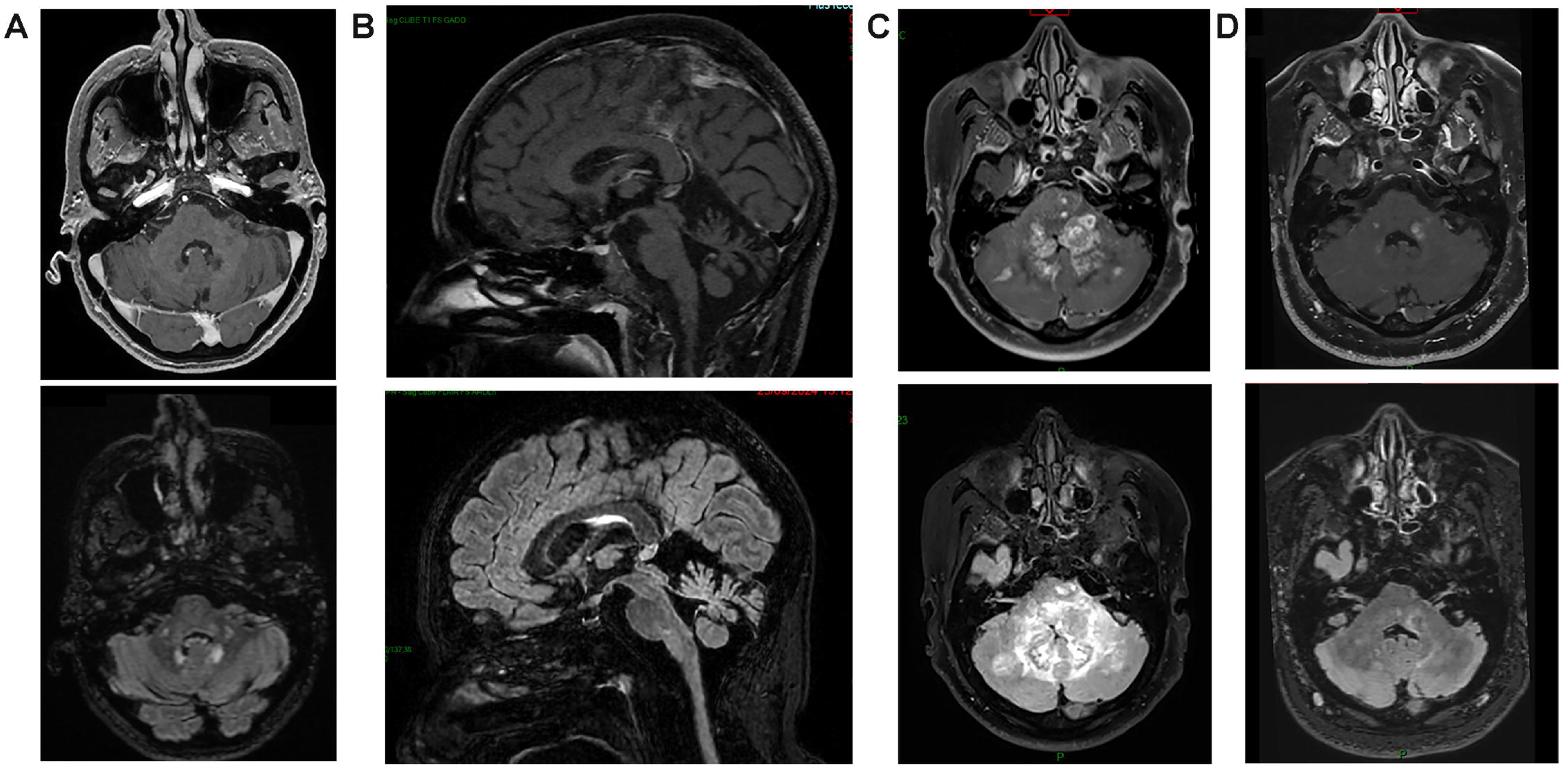
Brain Magnetic Resonance Imaging of Patients with BRAFV600E in Skin. A, B and C: top: T1-weighted images with contrast, bottom: T2-FLAIR weight image. Patient #042 with degenerative Erdheim Chester complicated by neurodegeneration: Abnormal high T2-FLAIR signal in the dentate nuclei area and in the cerebellar peduncles without contrast enhancement (top: T1-weighted images with contrast, bottom: T2-FLAIR weight image) (A); Atypical aspect of patient #062 (B) with major cerebellar atrophy occurring ten years after diagnosis of LCH; Atypical aspect of patient #020 with tumor-like infiltration on the posterior fossa (C); Response of these lesions after 3 months of treatment with dabrafenib and trametinib (D).

## DISCUSSION

The results from this prospective observational cohort study show that 85% of patients with histiocytosis harboring a *BRAF^V600E^* mutation also exhibit this mutation in healthy skin. When patients underwent multiple skin biopsies, *BRAF^V600E^* was found to be present in all. The presence of *BRAF^V600E^* in healthy skin is a micro-mosaicism and involves tissue-resident macrophages. Treatment with BRAF/MEK inhibitors, sometimes for longer than 5 years, had no effect on the variant allele frequency of *BRAF^V600E^*. Our findings also show that only patients with *BRAF^V600E^*-mutated healthy skin have neurodegenerative disease.

The detection the *BRAF^V600E^* in blood monocytes and bone marrow CD34+ progenitors^5,6^ supports the myeloid origin of Langerhans cell histiocytosis and Erdheim Chester disease. However, Langerhans cells and most resident macrophages of several organs, including microglia, do not usually derive from blood monocytes, but from yolk sac progenitors.^20^ Combining in-situ hybridization and immunohistochemistry with CD163 and CD4, we show that *BRAF^V600E^*-positive cells in healthy skin biopsies colocalize with dermal macrophages. Interestingly, 42% of the patients with *BRAF^V600E^*-positive skin did not have detectable *BRAF^V600E^* in blood or bone marrow cells. This excludes blood contamination of *BRAF^V600E^*-positive skin, and suggests that the *BRAF^V600E^* dermal macrophages of some patients may originate from progenitors of the yolk sac rather than from bone marrow.

Micro-mosaicism was recently demonstrated in the microglia of height /8 patients with *BRAF^V600E^* histiocytosis.^16^ We show here that 57/67 patients with *BRAF^V600E^* histiocytosis had a *BRAF^V600E^* micro-chimerism in skin. Mosaicism of *BRAF* oncogenic variants can be responsible for cardiofaciocutaneous syndrome, which is characterized by malformations, and is often associated with neurological manifestations.^21^ Mice with *BRAF^V600E^* micro-chimerism specifically targeted in macrophage lineage only develop late-onset neurodegeneration.^15^ Thus, clinical symptoms induced by micro-mosaicism in humans probably also depends on which cell lineage is involved, and our results show that *BRAF^V600E^* micro-chimerism in skin macrophages is associated with the occurrence of histiocytosis and/or neurodegeneration.

Most patients with multisystemic Langerhans cell histiocytosis have *BRAF^V600E^* in plasma cell free DNA.^22^ Surprisingly, *BRAF^V600E^* is usually still detectable in patients in clinical remission from Langerhans cell histiocytosis.^8^ We show here that long-term treatment of patients with BRAF/MEK inhibitors has no effect on variant allele frequency of *BRAF^V600E^* in skin, suggesting that *BRAF^V600E^*in plasma cell free DNA is produced by both tumor cells and resident macrophages.

The use of targeted therapies with BRAF or MEK inhibitors drastically changed the prognosis of patients with histiocytosis.^6,7^ However, neurodegeneration remains a major problem, because in most cases neurological symptoms only stabilize or partially regress with BRAF/MEK inhibitor treatment despite an otherwise good clinical response of tumor masses. It is thus extremely important to identify patients at risk of neurodegeneration and to detect neurodegeneration as early as possible. A systematic brain MRI is regularly performed during follow-up in children at high risk; however, subtle brain MRI abnormalities can remain undetected by radiologists who lack experience in this rare disease.

Several biomarkers are associated with neurodegeneration in patients with histiocytosis, including neurofilament light chain concentration in cerebrospinal fluid,^23^ hematologic involvement by histiocytosis,^24^ and detection of *BRAF^V600E^* in blood leukocytes.^25^ However, the low sensitivity of these markers does not drive prospective follow-up in these patients. In contrast, our study demonstrates a highly sensitive marker: all the patients without *BRAF^V600E^* in healthy skin were free of neurodegeneration. Thus, patients with histiocytosis but wild type skin could avoid follow-up with brain MRI. By contrast, 34% of patients with histiocytosis and *BRAF^V600E^* skin had neurodegeneration, although some were negative for *BRAF^V600E^* in blood cells or had minimally detectable changes on brain MRI. Whereas the molecular status of histiocytosis is still rarely investigated in patients, the skin *BRAF^V600E^* test does not require this previous analysis.

Patients can develop neurodegeneration when Langerhans cell histiocytosis has achieved remission, and when biopsies performed several years earlier are no longer available for molecular analysis. In our study, four patients whose molecular status was not available on histiocytosis were found to have *BRAF^V600E^* in skin samples. Because most bone lesions from Langerhans cell histiocytosis can regress spontaneously and others are asymptomatic, we suspect that some patients presenting with cerebellar ataxia may have had undiagnosed or asymptomatic Langerhans cell histiocytosis in the past. Thus, testing for *BRAF* mutations could be useful during the initial work-up of patients presenting with “idiopathic” cerebellar ataxia associated with diabetes insipidus, and/or patients with MRI findings suggestive of neurodegeneration related to Langerhans cell histiocytosis.

Currently there is no validated treatment for neurodegeneration in patients with histiocytosis; however, some patients treated with BRAF inhibitors can achieve stable disease and a minority show neurological responses.^26^ A mouse model also showed that early treatment with BRAF inhibitors delays symptoms of neurodegeneration.^15^ We show here that long-term treatment with BRAF/MEK, which was effective on tumors and inflammation, had no effect on the variant allele frequency of healthy skin. This lack of effect may also be the case in microglia, and thus could explain the lesser effects on neurodegeneration. Most histiocytosis cells can have oncogenic addiction and can be controlled by BRAF/MEK inhibitors, while most healthy *BRAF^V600E^*cells may only enter hibernation under a low concentration of drugs, and constitute a reservoir. Such a reservoir may be responsible for tumor relapse when treatment is stopped, and/or for microglia-induced neurodegeneration. Targeting the resident “healthy” histiocytes harboring *BRAF^V600E^* probably requires higher doses of BRAF inhibitors than targeting tumoral histiocytosis. Prospective trials including pharmacologic analyses should be performed in patients with neurodegeneration.

## ACKNOWLEDGMENTS

We thank Paul Takam Kamga, Claude Capron, Noémie Urvoy and Philippe Rameau for fruitful discussions and advices, and Espoir Anago, Mariama Bakari, Thamila Satour, Véronique Toulza and Neila Uteene for their excellent technical assistance. Sophie Rushton-Smith, Ph.D. (MedLink Healthcare Communications) provided editorial assistance, funded by Assistance Publique-Hôpitaux de Paris (AP-HP).

## AUTHORS’ CONTRIBUTIONS

JF Emile: Conceptualization, Data curation, Formal analysis, Funding acquisition, Investigation, Methodology, Project administration, Resources, Software Supervision, Validation, Visualization, Writing – original draft, review & editing, Z Hélias-Rodzewicz: Formal analysis, Investigation, Methodology, Software Supervision, Validation, Visualization, Writing – original draft, review & editing, M Papo, IA Ungureanu, C Baek, F Triboulet, TH Wong, R Ben Jannet, B Terrones, S Es-Qalli, PM Suret, B Reig, A Atallah, A Semenov, R Degrave, M Nguyen, R Sutera, E Siliogk, J de la Rochefoucauld, H Adle-Biassette, I Plu, A Mathian, C Catelain, J Razanamahery, C Ewenczyk, S Héritier, F Cohen-Aubart, P Hirsch, Z Amoura, J Donadieu: Investigation, Validation, and Writing – review & editing, A Idbaih: Investigation, Validation, Visualization, Writing – original draft, review & editing, J Haroche : Data curation, Formal analysis, Funding acquisition, Investigation, Project administration, Validation, Writing – original draft, review & editing

## ETHICS COMMITTEE APPROVAL

The study TARGET-HISTIO (ClinicalTrials.gov number, NCT04437381) was approved by the French Ethics and Scientific Committee for Health Research, Studies and Evaluations (CESREES 2814848 bis). Lumbar skin biopsy was considered minimally invasive according to French regulations.

## Data sharing statement

Data can be provided upon request to J.F. Emile.

